# Delineating the macroscale areal organization of the macaque cortex in vivo

**DOI:** 10.1101/155952

**Authors:** Ting Xu, Arnaud Falchier, Elinor L. Sullivan, Gary Linn, Julian Ramirez, Deborah Ross, Eric Feczko, Alexander Opitz, Jennifer Bagley, Darrick Sturgeon, Eric Earl, Oscar Miranda-Domínguez, Anders Perrone, R. Cameron Craddock, Charles Schroeder, Stan Colcombe, Damien Fair, Michael P. Milham

## Abstract

Complementing longstanding traditions centered around histology, fMRI approaches are rapidly maturing in delineating brain areal organization at the macroscale. The non-human primate (NHP) provides the opportunity to overcome critical barriers in translational research. Here, we establish the data and scanning conditions for achieving reproducible, stable and internally valid areal parcellations in individuals. We demonstrate that these functional boundaries serve as a functional fingerprint of the individual animals, and can be achieved under anesthesia or awake conditions (rest, naturalistic viewing), though differences between awake and anesthetized states precluded the detection of individual differences across states. Comparison of awake and anesthetized states suggested a more nuanced picture of changes in connectivity for higher order association areas, as well as visual and motor cortex. These results establish feasibility and data requirements for the generation of reproducible individual-specific parcellations in NHP, as well as provide insights into the impact of scan state and motivate efforts toward harmonizing protocols.

## INTRODUCTION

The non-human primate brain model remains one of the most promising vehicles for advancing translational neuroscience (Essen, 2012; Hutchison et al., 2015; Mars et al., 2011; Nelson and Winslow, 2008; Phillips et al., 2014; Rilling, 2014; Vanduffel et al., 2014). This point is particularly true for investigations requiring the definition or manipulation of specific cortical targets and their associated circuitry. Traditionally, such lines of inquiry have relied on histological techniques, which are capable of delineating specific cortical areas based on postmortem cytoarchitectonic properties. Beyond their labor intensiveness, such methodologies are inherently limited in their utility for the longitudinal examinations needed to study neuroplasticity, neuromodulation and brain development. The availability of histological atlases for various non-human primate species provides an alternative means of guiding intracranial measurements and interventions. However, such atlases are inherently limited in their ability to account for differences in brain areal organization among animals, which can negatively impact experimentation when not properly considered (Cerliani et al., 2017; Gordon et al., 2017a, 2017b; Van Essen et al., 2016).

Recent fMRI studies in the humans have demonstrated the feasibility of using resting state fMRI (R-fMRI) methodologies to delineate the functional brain organization of the cortex at the macroscale based on intrinsic brain function (Cohen et al., 2008; Craddock et al., 2011; Glasser et al., 2016a; Gordon et al., 2016; Wig et al., 2014). Not surprisingly, the postmortem histological parcellation approaches used to differentiate spatially contiguous areas into functional units based on differences in their cytoarchitectural and connectivity properties have served as a model for such efforts. In particular, Cohen et al. (2008) proposed a novel approach to perform areal parcellation in the human brain that measures gradients in intrinsic functional connectivity among neighboring vertices, which can, in turn, be used to identify transitions across distinct functional units. This approach has been extended to the full neocortex, with applications revealing functionally discrete areas at the group-average level that are homogenous and reproducible across studies (Glasser et al., 2016; Gordon et al., 2014; Schaefer et al., 2017). Several of the borders have been highlighted for their correspondence to task activations and cytoarchitectonically defined areas (Buckner and Yeo, 2014; Gordon et al., 2017a; Wig et al., 2014), though such relations are yet to be fully validated at the individual level. Most recently, this gradient-based boundary mapping approach has been applied at the individual-level in human, successfully mapping full-brain gradient and transition patterns that despite commonalities appear to be individual-specific (Gordon et al., 2017a, 2017b; Laumann et al., 2015; Xu et al., 2016) and have moderate to high test-retest reliability using as little as 20 minutes of data.

Work by several groups has suggested the possibility of parcellating the macaque brain in-vivo as well, using methods such as functional MRI, diffusion tractography and invasive neural recordings. However, to date, these studies (Mars et al., 2011; Neubert et al., 2015; Schönwiesner et al., 2014; Vanduffel et al., 2014b) have primarily focused on more limited sections of cortex, e.g. visual cortex, including occipito-temporal, parietal and frontal cortex. The logical extension of this work – fMRI-based parcellation of the neocortex of the macaque in its entirety – would be extremely valuable in evaluating the strengths and limitations of the macaque as a translational model (Hutchison and Everling, 2012, 2014; Hutchison et al., 2012a, 2012b; Miranda-Domínguez et al., 2014; Shen et al., 2012).

In the present work, we pursue this aim using gradient-based fMRI parcellation methods to arrive at a putative map of the functional brain organization of the macaque cortex in its entirety at individual level. Although promising in human literature, there are several challenges that need to be addressed to achieve this goal, namely, 1) the feasibility of the boundary mapping parcellation in an individual macaque; 2) the validity of the parcellation; 3) the reliability of the gradient-based boundary mapping approach across scans in an individual macaque; and 4) the stability of the parcellation and the quantity of individual data required for an accurate estimate of full-brain functional areal organization. To address these challenges, we made use of four adult rhesus macaque datasets from two independent research institutions: Nathan Kline Institute (NKI-Dataset; n=2) and Oregon Health and Science University (OHSU-Dataset; n=2). Each macaque dataset includes between 294 and 688 min of awake and/or anesthetized fMRI scans. Using these data, we first identified the individual-specific functional boundaries and areal parcellation for each dataset and compared it against the available anatomical borders from previous studies. Then, we assessed the functional homogeneity of the gradient-based areal parcellation and demonstrated the uniqueness of the result obtained for each specific monkey. We also evaluated the stability of functional connectivity (FC) profiles, gradient and boundary maps to investigate the total quantity of data required for estimating full-brain functional areal organization. In addition, we examined the impact of a monocrystalline iron oxide ferumoxytol (MION) contrast agent, on the quality of data needs for parcellation. As prior work has demonstrated its utility in increasing the ability to detect functional connectivity patterns for non-human primates, when scanning at 3.0T (Grayson et al., 2016). Similarly, we examined the potential influences of anesthesia, which is known to decrease measures of FC somewhat (Hutchison et al., 2014a). Together, these analyses establish feasibility and data requirements for the generation of reproducible individual-specific parcellations in the non-human primate, as well provide insights into the impact of varying factors (e.g., scan state) on findings.

## RESULTS

For each individual macaque, the gradient-based parcellations were calculated for each different condition (awake or anesthesia) and contrast agent status (with or without MION). The gradient evaluates the spatial transitions in the similarity of neighboring FC profiles across the native cortical surface. Corresponding edges and parcels across animals were identified from the gradient maps using the ‘watershed’ algorithm (Gordon et al., 2016). The final functional boundary (i.e. edge density) were averaged across cortical surface which represented the possibility that a vertex was identified as an edge. The details of this method were described in previous studies (Laumann et al., 2015; Xu et al., 2016) and a brief description of the method is presented in the methods. Of note, the awake fMRI data were acquired with naturalistic viewing paradigm – movies were playing during the awake scans for the NKI dataset (monkey ID: NKI-R, NKI-W). To investigate the impact of differing awake states (movie and rest), we additionally collected one pure rest session for one animal (NKI-W). In the following section, ‘awake’ states without any specifications refer to naturalistic viewing states.

### High Reproducibility of Gradient-based Parcellations

We first focused on the awake states with MION contrast collected from the NKI site (Figure 1). Within each individual animal, the gradients and edge density detected for the awake state with MION demonstrated high reproducibility across individual scans (see all animals in supplement Figure S2). Consistent with prior work in humans (Xu et al., 2016; Gordon et al., 2017), gradient maps showed a higher degree of reproducibility than edge maps when equivalent amount of data were used. In order to obtain a better assessment of the upper bound for reproducibility, we randomly divided the sessions for each macaque into two subsets (details see Supplement Table S1) and regenerated gradient and edge maps for the data available in each subset. Markedly higher reproducibility was shown for each of the measures, as well as the resultant parcellation maps (see Figure 1 for edge overlap map). The spatial correlations between two gradient maps for macaque NKI-R and NKI-W while awake were 0.71 and 0.81; for edge density, the correlations were 0.30 and 0.47. The Dice coefficient between two parcellations were 0.68 and 0.71.

**Figure 1:**
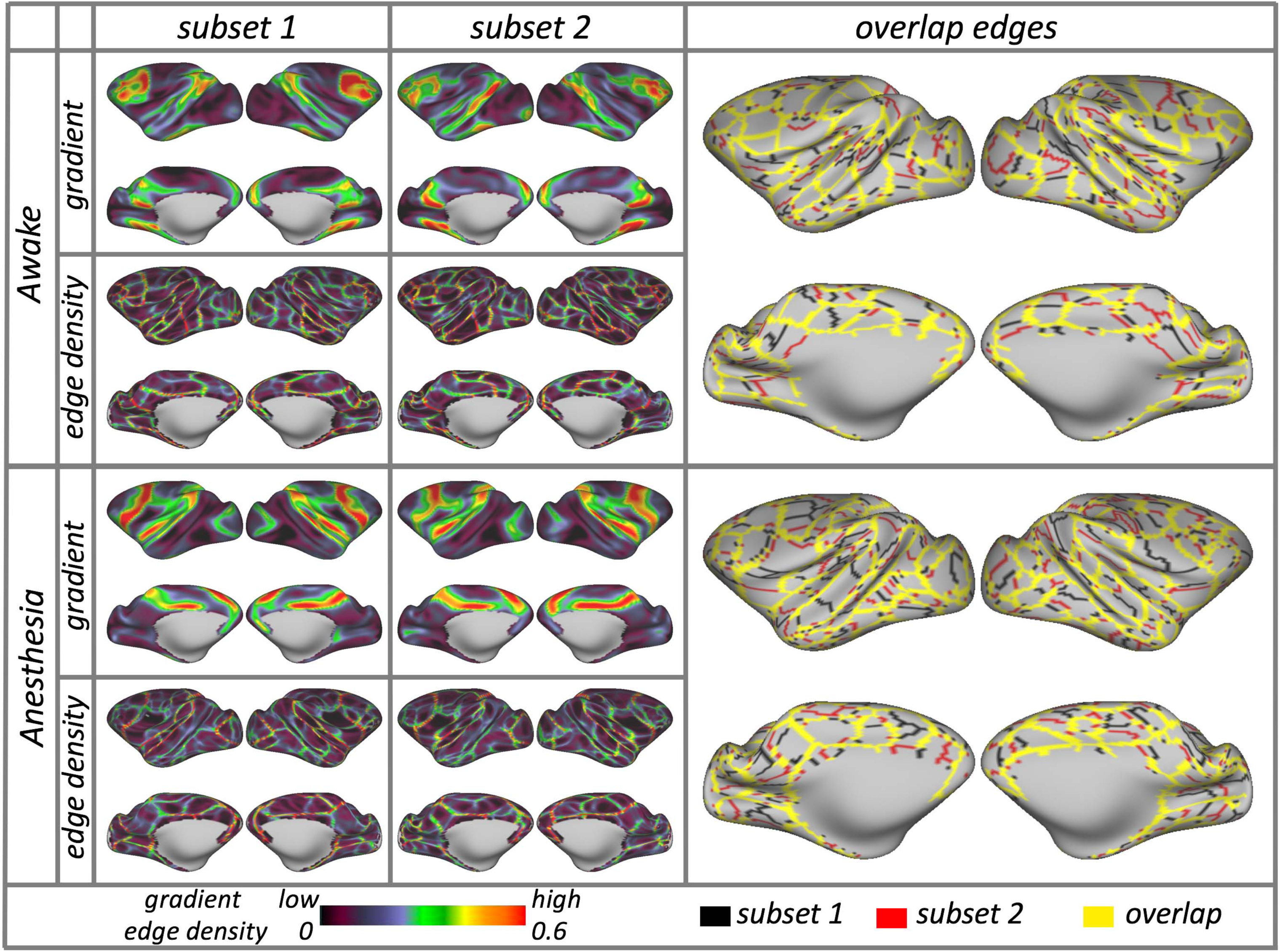
Subject-specific areal organization is highly reproducible across two independent subsets of sessions under awake and anesthesia (examples of gradient, edge density and overlapped edges in NKI-W, see Figure S3 for all animals).

Next we looked at data obtained during the anesthetized state using MION in subjects from each site, NKI and OHSU. Similar to the awake results, we observed a high degree of reproducibility in gradients, edge density and corresponding parcellations (see representative subjects in Figure 1-lower and all subjects in Figure S1). The spatial correlations in gradients between subsets were 0.79 (NKI-R), 0.87 (NKI-W), 0.84 (OHSU-1), and 0.90 (OHSU-2); for the edge density were 0.30, 0.48, 0.57 and 0.52.

The Dice coefficient between parcellations were 0.69, 0.70, 0.70, and 0.72, respectively.

In addition, we examined the reproducibility of gradient, edge density and parcellations using data obtained without MION (i.e. BOLD). Compared with the data using MION, the findings were similar but less reproducible. Specifically, in the two awake NKI datasets (NKI-R, NKI-W), the spatial correlations between two subsets were 0.32 and 0.54 for the gradient maps, and 0.20 and 0.34 for the edge density maps; the Dice coefficients were 0.67 and 0.66 for the final parcellations. In the anesthetized data from OHSU, the spatial correlations were 0.69 and 0.78 in gradients, and 0.65 and 0.59 in edge density. The Dice coefficient were 0.70 and 0.67 for the final parcellations.

### Parcellation Homogeneity

Consistent with prior work in humans (Lauman et al., 2016), we assessed the internal validity of the parcellation maps obtained for each macaque. Specifically, we calculated the homogeneity of FC similarity in the parcels generated from subset 2 (reference subset) using the data from subset 1 (test subset). The mean homogeneity across all parcels in the awake with MION condition was 0.91 for NKI-W and 0.77 for NKI-R, while under anesthesia across the four macaques was range from 0.85 to 0.90 (mean=0.88 SD=0.019, see Figure 2, red line). Then, we assessed the degree to which this parcellation was more homogenous than a null distribution of mean homogeneities generated from 1000 ‘random’ parcellation maps for the same subject; each of these maps were generated by randomly rotating the parcellation units around the cortical surface for each subject (Gordon et al., 2016). The mean homogeneity of null model parcellations was range from 0.73 to 0.86 (mean=0.82, SD=0.048) for four macaques (these findings are comparable to those previously reported in humans (Laumann et al., 2015; Xu et al., 2016); in each macaque, the real parcellations were significantly greater than that obtained in null model parcellations (awake: Z score = 5.94, 10.79, p<0.001; anesthesia: 9.20, 9.56, 9.00, 7.22, p<0.001, Figure 2).

**Figure 2:**
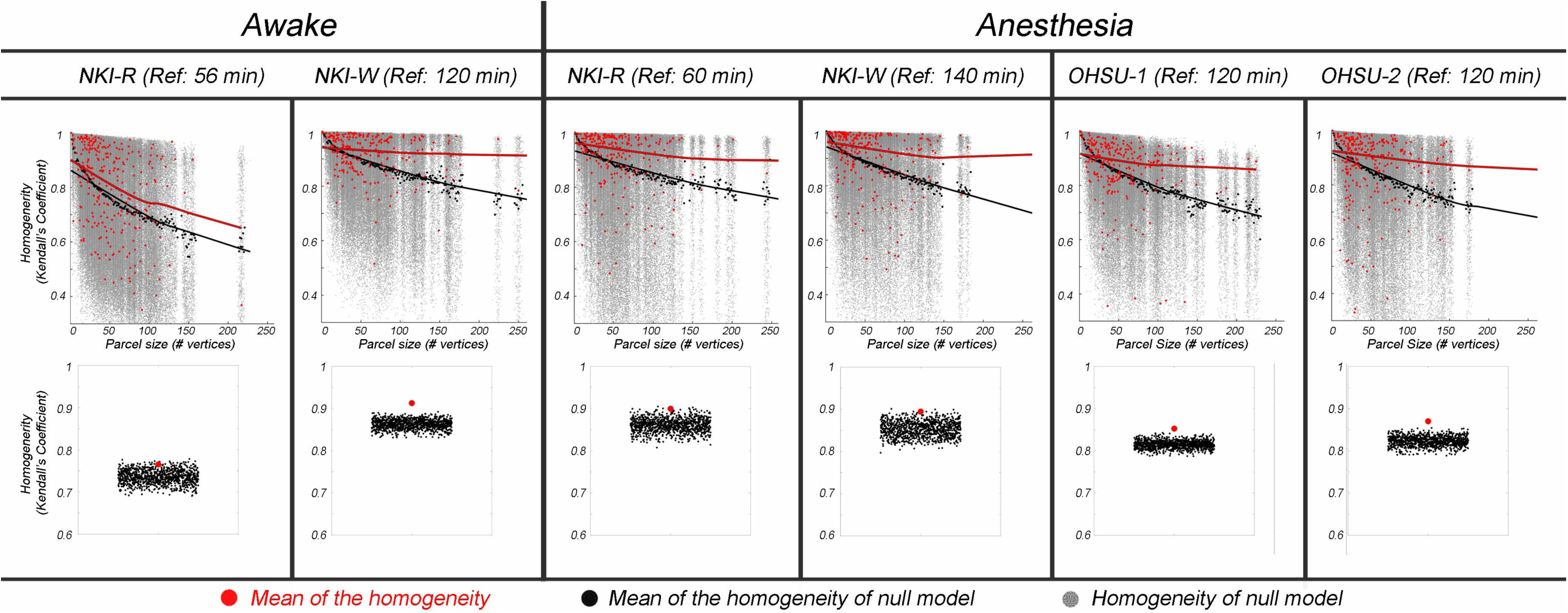
Upper panel: Homogeneity of real parcels (red dots) by parcel size compared to homogeneity of null model parcels (gray dots) under awake and anesthesia states. The real homogeneities were calculated in the parcels generated form subset 2 (reference subset) using the data from subset 1. Black dots were the mean homogeneity across iterations for each null model parcel. The lines are the LOWESS (locally weighted scatterplot smoothing) fit represent the effect of parcel size on homogeneity of the subject parcels (red line) and the null model parcels (black line). Lower panel: The mean homogeneity across real parcels (red dot) for each monkey under awake and anesthesia states are significantly higher than the mean homogeneity from the null model parcellations (black dots).

We noted that the parcellation from the awake condition, with MION, for macaque NKI-W was more homogeneous than NKI-R. This discrepancy arose partially because of the total amount of data used for NKI-W, which was about three times that for NKI-R. Accordingly, we randomly sampled subset 1 and examined the relationship between the homogeneity and amount of scan time. As we expected, the homogeneity increased with the scan time (Figure 3, the Lowess fit line plotted the homogeneity at each increment of data used). Consistent with expectations based on prior work in humans, the homogeneity rapidly increased with 30 min of data (average incensement = 0.09), with relatively modest increases as additional data is added (Figure 3).

**Figure 3:**
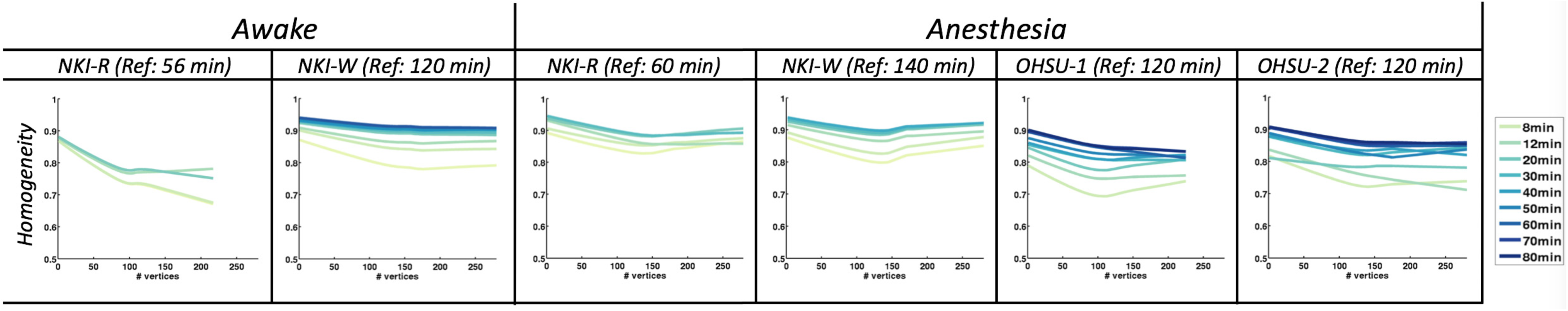
The homogeneity of parcellations vary with the amount of test data. The Lowess fit line plotted the homogeneity at each increment of data used in subset 1 under same condition for each macaque.

Using the same test subset (subset 1) as above, we also used homogeneity to compare the fit of the individual-specific, functionally-derived parcellations in the present work, with those of five atlas-based parcellations for the macaque (LV00: Lewis and Van Essen 2000; Markov11: Markov et al., 2011; PHT00: Paxinos and Franklin 2000; FV91: Felleman and Van Essen 1991; B05: Brodmann 1905). These five atlases were mostely defined upon architectonics in a single animal (LV00 was based on 5 specimens). Van Essen’s group have registered the those parcellation to F99 macaque surface enable us to compare the homogeneity with the functional-derived parcellation here (Van Essen et al., 2012). Specifically, we compared the homogeneities of each alternative atlas-parcellation against the homogeneities of a null model (based on that specific parcellation), which was generated using the same procedure as above (i.e. 1000 random rotations of the parcellations units around the cortical surface). The individual-specific, functionally derived parcellations (based on the data from subset 2), were more homogenous than any of the alternative parcellations (Figure 4, upper). Across the five atlases, only LV00 showed consistently greater homogeneity (than the null model) for both NKI-R and NKI-W, and that was limited to the awake condition with MION (Figure 4, lower). The PHT00, FV91, Markov11, B05 were not significantly more homogenous than their null models in the awake condition for either, NKI-R or NKI-W. In the anesthetized state, all five atlases failed to show significantly greater homogenous than their corresponding null models in at least 3 of the 4 macaques (NKI-W, OHSU-1, OHSU-2). Only PHT00, LV00, Markov11, B05 were significantly better than their corresponding null models in 1 animal (NKI-R).

**Figure 4:**
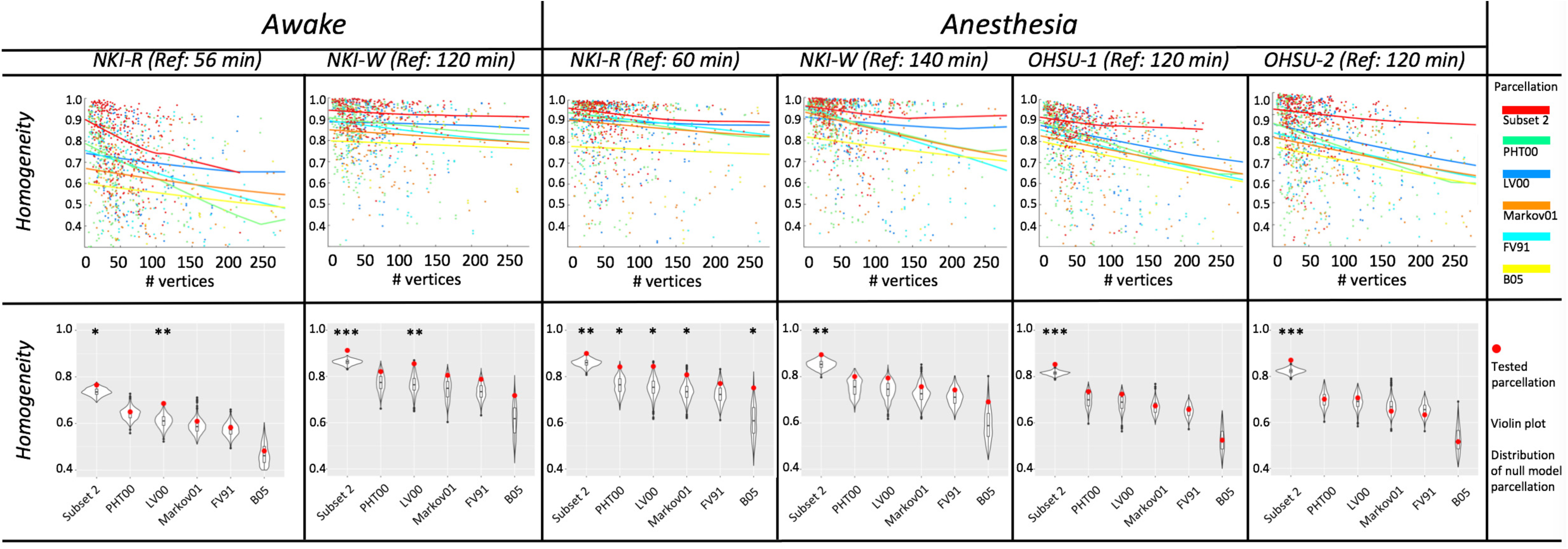
The homogeneity of functional-defined parcellation is more homogenous than any other parcellation under awake and anesthesia states. Top: homogeneities tested in subset 1 at each of parcel size for functional-defined parcellation (red line) and five atlases. Bottom: weighted mean homogeneity across parcels of functional-defined parcellations and atlases (red dots) compared with the averaged homogeneity across parcels of each of null model (distribution of 1000 randomization in violin plot). ***indicates p<0.001, ** indicates p<0.01, * indicates p<0.05 in its 1000 null model randomizations.

Additionally, we used homogeneity to confirm that the findings from functionally-derived parcellations exhibited individual specificity. Specifically, we compared the homogeneity of an appropriately matched parcellation (i.e., homogeneity calculation and parcellation based on two non-overlapping data subsets from the same subject) vs. inappropriately matched parcellation (i.e., homogeneity calculation and parcellation based on data subsets from different subjects). As part of this examination, we also examined the specificity of findings for an individual with respect to the scan condition and contrast agent status). In 17 of 20 subsets, we found homogeneity to be maximal when the individual, scan condition and contrast agent status were all appropriately match (red triangles in Figure 5) versus not (grey dots in Figure 5; results from the five atlases are depicted in blue).

**Figure 5:**
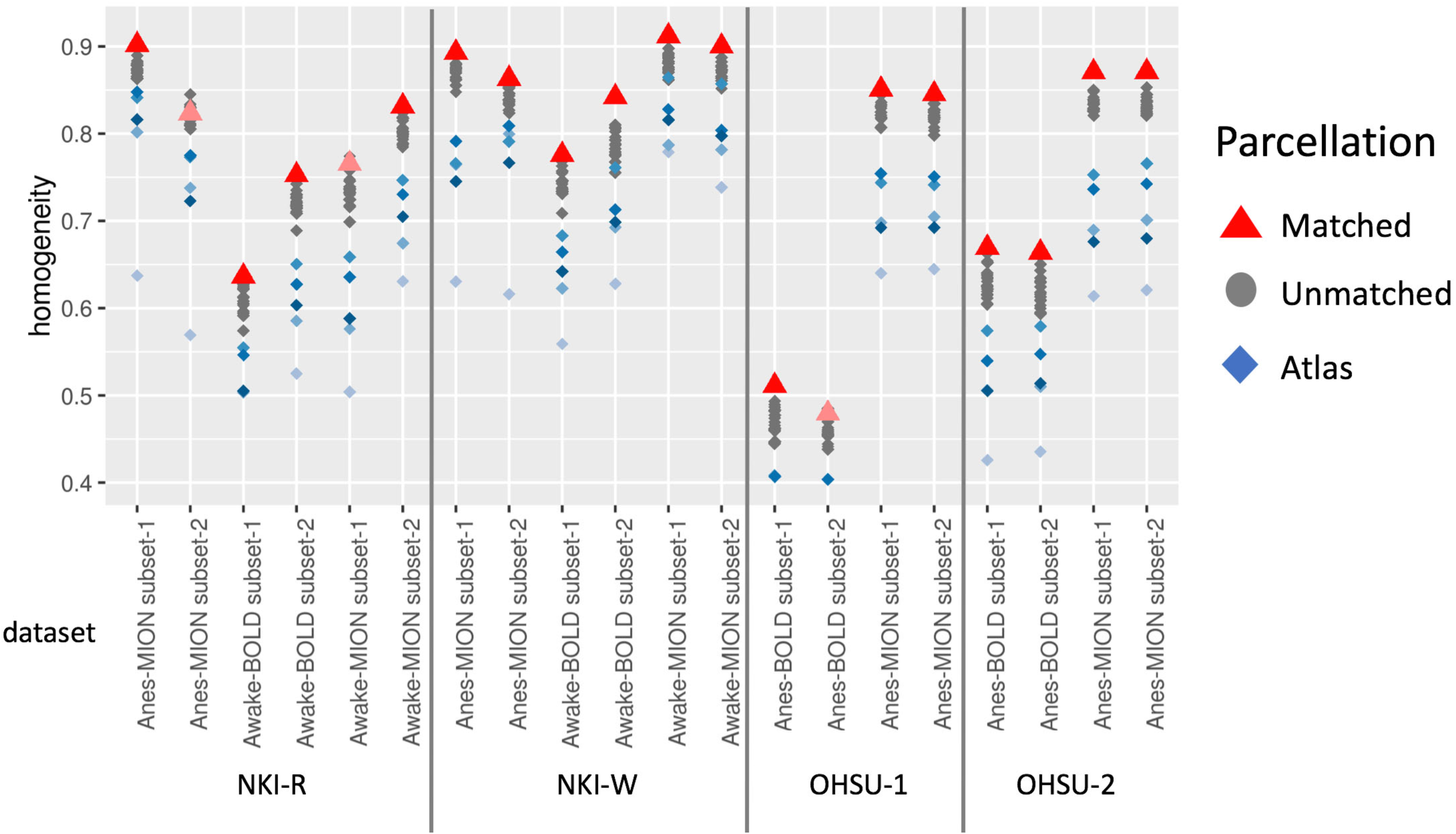
The average homogeneity was higher within individual specific functional-derived parcellation under same condition (red triangle) than inappropriately matched parcellation (gray dots indicate the homogeneity calculation and parcellation based on different condition or subjects; Blue dots indicate the average homogeneity based on five atlases).

We noted that the homogeneity scores based on BOLD conditions dataset were lower than MION conditions within each macaque. Thus, we compared the homogeneity across different states (awake vs. anesthesia) and contrast types (MION vs. BOLD). The MION data exhibited higher homogeneity than BOLD condition regardless of awake (Figure S4, blue line) or anesthetized states (Figure S4, turquoise line); the z scores between MION and BOLD data (t-test) in awake state are 8.94 (NKI-R), 11.14 (NKI-W), both p<0.001, while under anesthesia are 22.16 (OHSU-1) and 14.12 (OHSU-2), both p<0.001. These differences between awake and anesthesia are examined and discussed in the later sections.

### Comparison of Parcellation with Known Topographic Areas

To further assess the validity of the parcellation maps using an external reference, the putative boundaries delineated by our functional gradient-based parcellation was overlaid with the areas 3, 4 and 17 borders previously established using post-mortem histology (Brodmann, 1909; Van Essen and Dierker, 2007). As depicted in Figure 6, the parcels derived by gradient-based mapping approximately overlapped with areas 3, 4 and 17 borders. This suggests that for these specific areas, gradient-based boundaries appeared to represent differences in function that are captured by histologic and topographic mapping.

**Figure 6:**
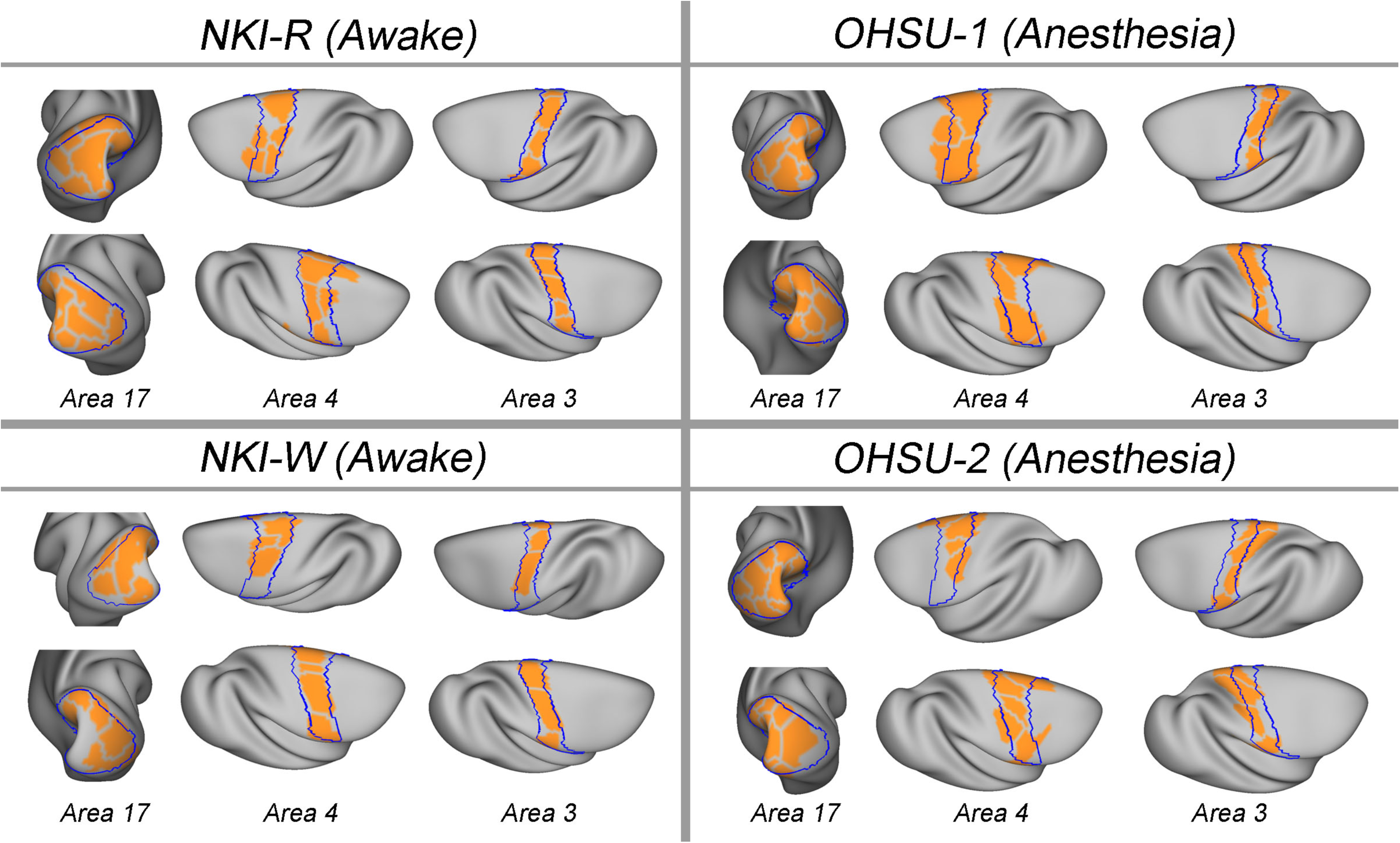
Subject-specific parcels (orange) overlaid on boundaries (blue) of post-mortem histologically defined areas (area 17, area 3, and area 4) from Brodmann (1905).

It is important to note that while there were some clear correspondences between our findings and those of the postmortem map obtained from prior work, some notable differences exist. While such differences likely suggest either greater detectability or the detection of a unique aspect of the functional architecture in the macaque, the possibility of artifactual findings related to the imaging also need to be considered in some cases. For example, in early visual cortices (e.g., the fractionates in area 17), we found additional edges not apparent in the postmortem map; examination of the raw data for these divisions suggests they may reflect an artifact arising from incomplete coverage, though alternatively, they may reflect the presence of further subdivisions. Future work with optimized imaging will be required to help differentiate between these possibilities.

### Comparison of Parcellation with fMRI Task Activations (Somatosensory Stimulation)

To provide additional insights into the relative fit of our functionally-derived parcellations with those of individual atlases. In this regard, we took advantage of the availability of somatosensory stimulation data obtained for NKI-R and NKI-W using data obtained during BOLD awake scanning. Figure 7 depicts left hemisphere activation during right hand stimulation with the parcellation overlaid on top of it. Consistent with prior work in humans, visual examination of activations and deactivations associated with stimulation appears to suggest that in many cases functional activations and deactivations appear to be at least grossly bound by the functional brain organization revealed by parcellation. To further test this point, we compared the fit of activations obtained for each of the animals with their respective parcellations and those of the five atlases. The inhomogeneity was calculated to access the fitness by computing the weighted standard deviation of z-values from the task activation for each parcel (Schaefer et al., 2017). A lower inhomogeneity of a parcellation indicates higher fitness of a given parcellation and activation. In both NKI-R and NKI-W, the parcellation driven by the condition matched functional data (awake BOLD) had better correspondence than atlases.

**Figure 7:**
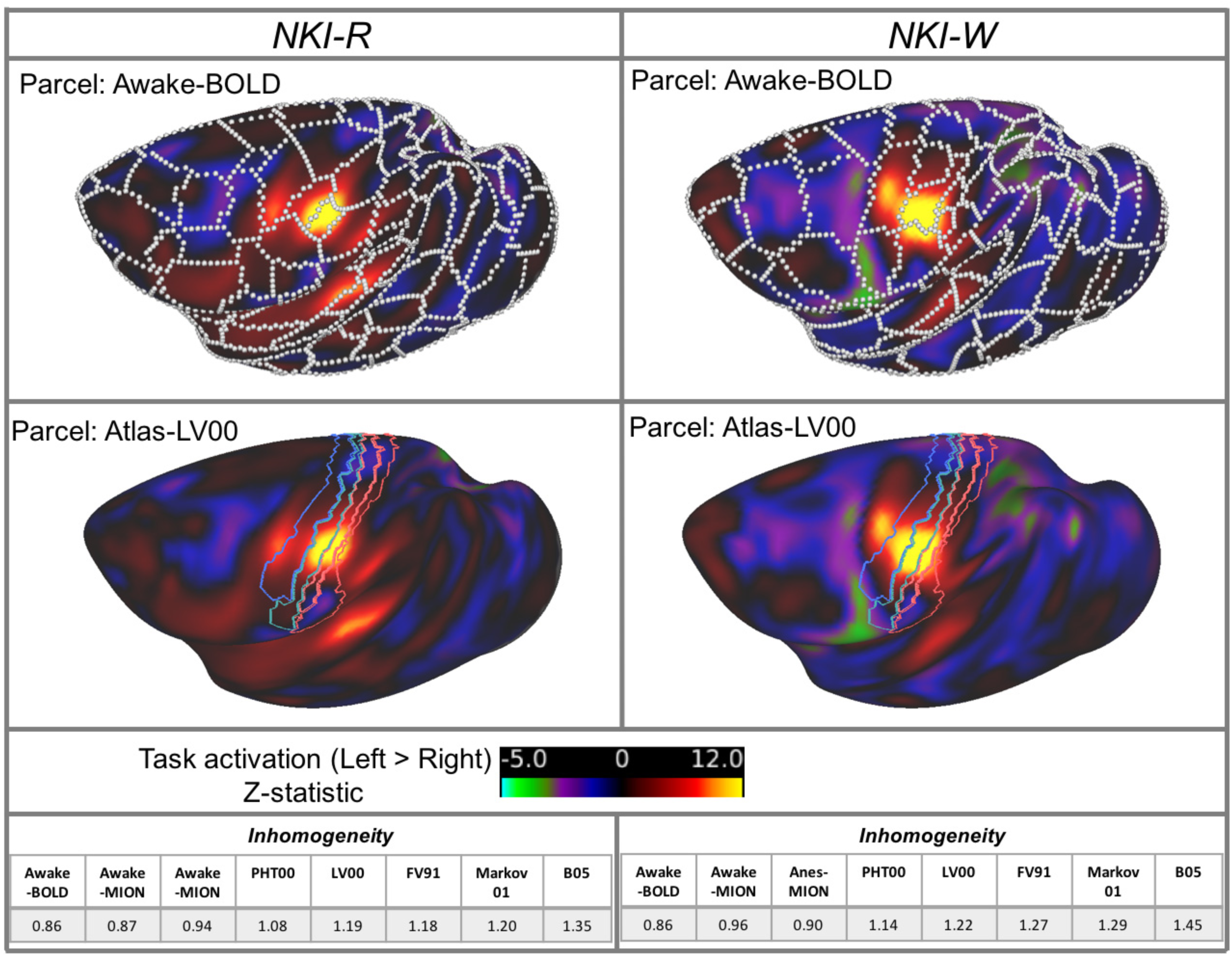
Task activations are correspondence with individual- and condition-specific functional defined boundaries. Z-statistic map of left > right contrast of BOLD condition for NKI-R and NKI-W overlapped with the edges (white borders) defined by naturalistic viewing BOLD sessions of same animal (upper panel). Z-statistic map of left > right contrast of BOLD condition for NKI-R and NKI-W overlapped with sensorimotor areas defined by atlas-PHT00.

### Data Requirements for Mapping the Reliable FC, Gradient and Edges

A key challenge for imaging efforts focused on brain parcellation is the determination of minimal data requirements to reliably and consistently capture individual parcellations. While the human imaging literature is actively working to establish data requirements, functional MRI imaging in non-human primates faces additional challenges. In particular, decreased signal to noise characteristics relative to humans due to smaller voxels size required, and the usage of anesthetics. The iron-based contrast agent, MION, is increasingly being used to overcome these challenges, particularly when scanning at 3.0T (Gautama et al., 2003; Grayson et al., 2016; Leite et al., 2002); however, the necessity of MION, particularly in the awake macaque, is not clear.

Here, we first examined the stability of FC, gradient and edge maps derived from data obtained during the awake state using MION in the NKI-dataset. More specifically, for each of the two macaques imaged in the awake state, we randomly split the data obtained with MION into two independent subsets (see table S1 for the details) and used subset 2 to derive reference FC, gradients and edge density. Next, we randomly sampled subset 1, started at 8-minute amount, adding in six 4-min increments, followed by three 8-minute increments of data, and then (for subject NKI-W only) two 16 minute increments (i.e. 8, 12, 16, 20, 24, 28, 32, 40, 48, 56, 72, 88 min); the maps generated from each increment were compared to the reference maps in the same macaque. The results convergent estimation for FC are averaged from 1,000 random samplings of data at each time point. Of note, to decrease the computational costs of gradient and edge density calculation for the random samples at each time point, we only used 100 random samples and based our calculation on a reduced (i.e., downsampled) set of vertices (i.e., 400) (prior testing found the results obtained to be comparable with those obtained for the higher resolution representations). The spatial correlation (Pearson’s r) was used to examine the similarity of FC and gradient while Dice coefficient was for measuring the similarity of the parcellations between test and reference dataset.

Consistent with prior human work, the average FC, gradients and parcellation from awake state (Figure 8, green line) were progressively increased by the amount of data incorporated into the analysis. Focusing on the subject NKI-W, who has more data, the average spatial correlation observed for 8 min from subset 1 with subset 2 (reference) was 0.80 (SD=0.063) for FC, and 0.56 (SD=0.122) for gradient maps; the mean Dice coefficient was 0.68 (SD=0.017) for the parcellations. The correlations were rapidly increased to 0.90 (SD=0.027) for FC and 0.70 (SD=0.071) for gradients with 28 minutes of data; mean Dice coefficient increased to 0.70 (SD=0.016) for parcellations. The averaged correlations increased more slowly up to 88 minutes, where they were 0.92 (SD=0.007) for FC, and r=0.74 (SD=0.009) for gradients; the Dice coefficient was 0.72 (SD=0.009) for parcellations with 88 min of data. Similar results were found in subject NKI-R, though the overall similarities were lower than subject NKI-W as less data was collected.

**Figure 8:**
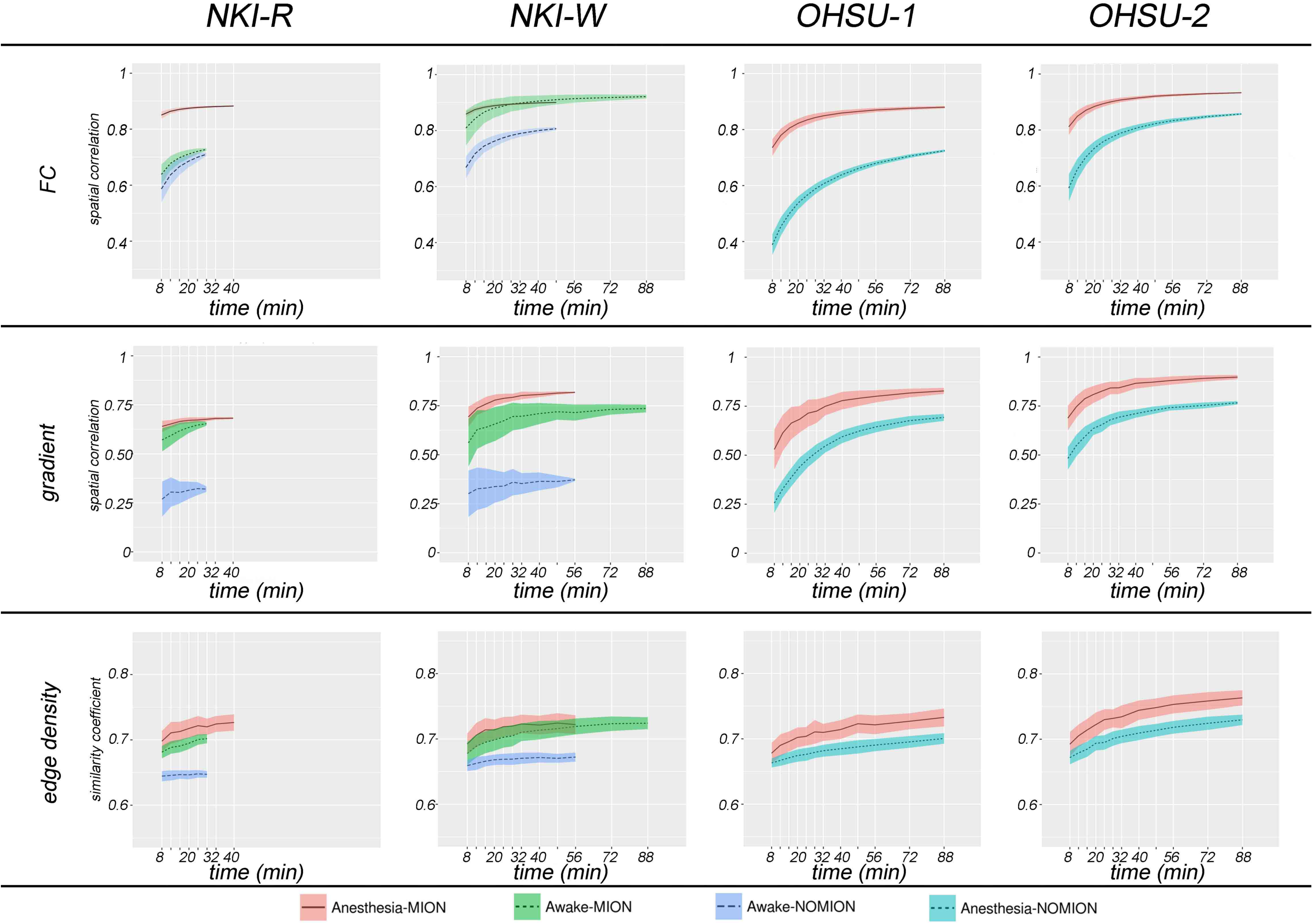
The stability of subject-specific functional connectivity, gradient, and edge density as a function of the amount of data used within different conditions (awake vs. anesthesia, with MION vs. without MION). Each line represents the mean (black line) and SD (colored patch) of similarity within each condition for each monkey; the similarity between maps generated from each incremental amount of data (from subset 1) and those from the other half of data (reference: subset 2).

Next we looked at data obtained during the anesthetized state using MION in subjects from each site, NKI and OHSU. Again, we compared findings obtained using two distinct subsets of data in each subject. We found that the stability profiles were highly similar to those observed in the awake state (Figure 8, red line). For each of the monkeys, the parcel-based FC, gradient and corresponding parcellations uninterruptedly increased from 8 min to 28 min and began to plateau after 40~56 min. At the 8 min point, the average similarities (spatial correlation) across four monkey from subset 1 with subset 2 (reference) were only 0.77 (SD=0.388) for FC and 0.61 (SD=0.310) for gradient; mean Dice coefficient was 0.69 (SD=0.020) for parcellations, while increased to 0.92 (SD=0.037) for FC, 0.83 (SD=0.343) for gradient, and 0.73 (SD=0.365) for parcellations at 40 min. The similarity is increased to 0.91 (SD=0.026) for FC, 0.87 (SD=0.033) for gradient, and 0.75 (SD=0.014) for parcellation at 110 min in the OHSU dataset.

In order to establish the impact of MION, for each of the four macaques, we also examined the stability of FC, gradient, edge density generated using the data obtained without MION (Figure 8, awake: blue line, anesthesia: turquoise line). When the data obtained for each monkey without the use of MION was divided into two subsets, the correspondence remains notably lower than obtained with MION - regardless of whether anesthesia was used (OHSU-1, OHSU-2) or not (NKI-W, NK-R).

### Identification of Unique Individual Areal Boundaries (Fingerprinting)

Next, we worked to confirm that despite apparent commonalities, the parcellations obtained were unique to each individual. To accomplish this, we assessed the similarity of the gradients and edge density generated across the four macaques and subsets (i.e. subset 1 and subset 2 for each macaque); Figure S5 depicts the spatial correlations between 21 gradient maps (2 macaque x 2 subsets x 3 conditions from NKI and 2 macaques x 2 subsets x 2 conditions from OHSU), as well as between 21 edge density maps (Figure S5). To facilitate the visualization, we highlighted the spatial correlations from MION data in Figure 9. The correlations between subsets within a given subject in the same state (awake or anesthetized - red and dark red dots, respectively), are notably greater than those between different subjects (blue and turquoise dots). When looking within the same state, an individual can be successfully identified using half of dataset for each macaque. Of note, the correlations in gradient and edge density across the monkey while under anesthesia (cadet blue dots) is higher than while awake (turquoise dots) - possibly suggesting that functional boundaries are more prone to variation while awake. Across states, measures obtained from data in the same macaque under two different awake states (i.e., naturalistic viewing, rest) exhibited a relatively high degree of similarity (fuchsia dots). However, the correlations in findings between awake and anesthetized (yellow dots) states in the same macaque were much less than those obtained in the same state. In addition, we also observed site effects, with correlations between monkeys across two sites (purple dots) being lower than within the same site while under the same circumstances (e.g., Anesthesia with MION).

**Figure 9:**
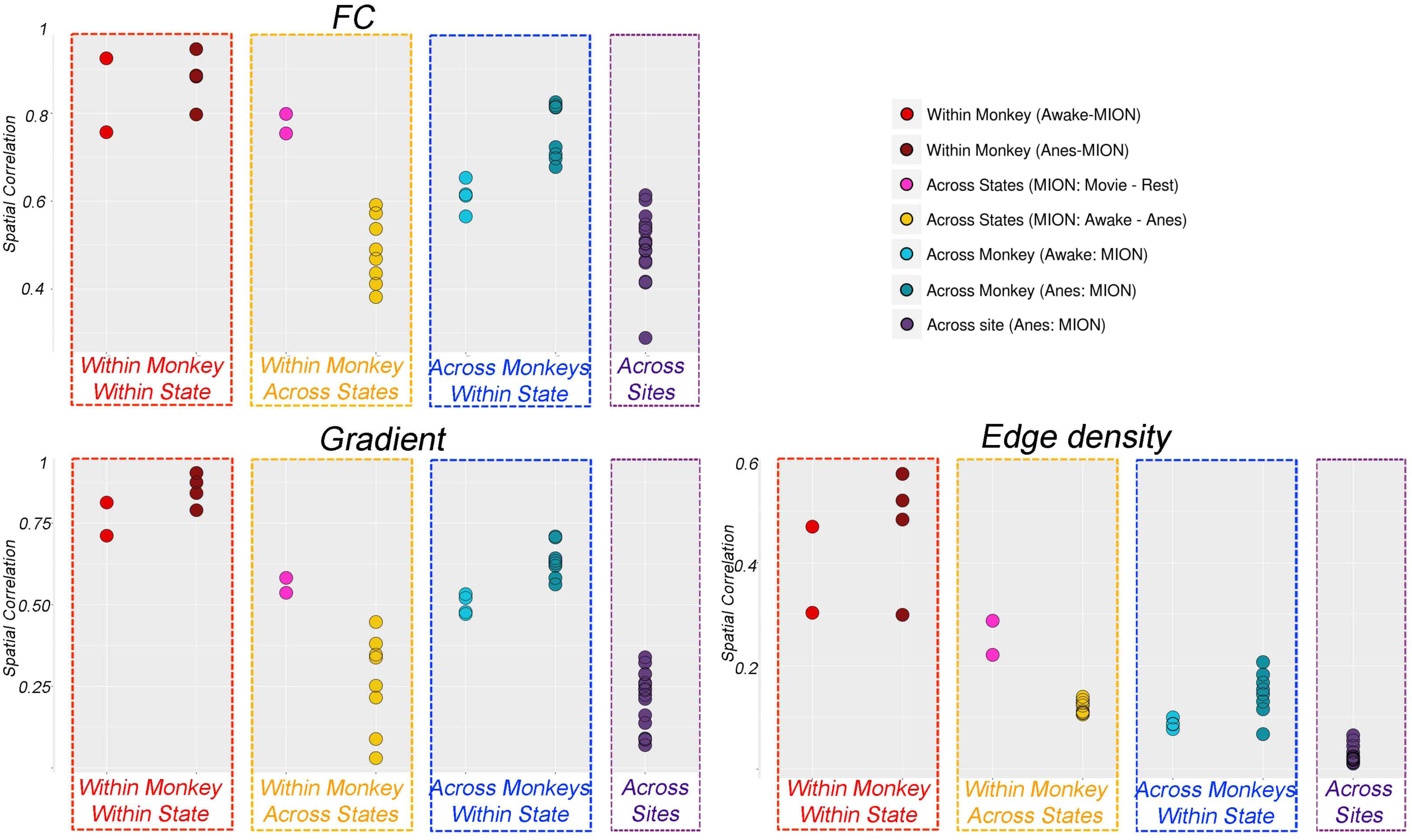
Dots plots of the spatial correlations in FC, gradient and edge density from two different subsets (subset 1 and subset 2), three conditions (awake movie, awake rest, anesthesia), and four monkeys from two sites. To facilitate comparison, we mark off dots to highlight the effects of subset, state, individual monkey and scan site in colored boxes.

### Effect of MION

As can be seen in Figure S5 (right panel), compared to the MION data within the same macaque, the FC, gradient and edge density maps obtained without MION have a relatively modest correspondence with those obtained using MION in anesthesia data (dark green dots). Similar but relatively lower correspondences were observed in the awake data from NKI-dataset (light green dots). These findings suggest that the signal obtained with MION is notably improved relatively to standard BOLD fMRI.

### Effect of State During Awake Imaging

Here, we examined the impact of differences among awake states by comparing functional connectivity, gradient and edge maps produced during the viewing of movies with those obtained during rest. Human imaging studies have drawn attention to the overall stability of functional connectivity patterns across differing awake conditions, despite more fine-grained modifications in connectivity associated with manipulations of state (e.g., rest, movies, task performance). Consistent with the findings from humans, we found a high degree of similarity between functional connectivity patterns (r=0.79) associated with rest and those observed with movie viewing. At the network level, a similar connection profile was found within and between networks except lateral visual cortex, which showed greater within network connectivity in movie viewing than rest. We also found a relatively high similarity for each, the gradient maps and the parcellations, generated using the two awaken states (movie and rest). The spatial correlation of gradient was 0.59 and Dice coefficient of parcellations was 0.68. As the similarity of FC, gradient and edge density between two awake states (movie and rest) within each monkey (see Figure 9 fuchsia dots) is higher than across monkeys in the awake movie state, this suggests that naturalistic viewing, which is more favorably tolerated by non-human primates, can be used as a flexible paradigm for awake imaging.

### Effect of Anesthesia

As reported in the prior section, the FC, gradient and edge density generated from differing subsets of data obtained with anesthesia showed a high degree of correspondence, similar to what was observed when subsets of data collected while awake were compared. However, the correspondence of areal organization measures was substantially lower when compared across the awake and anesthetized states in the same subjects (NKI-R, NKI-W). Specifically, compared to the awake condition, the spatial correlations in FC between awake and anesthesia was 0.52 (SD=0.08) (Figure 9 top panel, yellow dots). The gradient and edge density showed even less similarity between awake and anesthetized, with the spatial correlations across states with half subsets of data in NKI-dataset being 0.25 (SD=0.14) and 0.12 (SD=0.02) respectively (Figure 9, lower panel, yellow dots).

The lower correspondence of findings between states (awake, anesthetized) relative to within a state was not necessarily surprising. Previous studies have suggested that the spatial and temporal properties of functional connectivity observed in non-human primates can be impacted by anesthesia, with the profundity of the efforts observed depending on the depth of anesthesia (Hutchison et al., 2012; 2013; Vincent 2007). Visual examination of the FC matrices for the awake states (rest, movies) and anesthesia revealed notable regional connectivity differences that appeared to be specific to anesthesia (Figure 10B). To facilitate detection of those connections with the greatest change, for each monkey, we: 1) calculated the difference score (FC[awake] -FC[anesthesia]) for each connection, and then 2) transformed them to Z-scores based upon the mean and standard deviation for difference scores throughout the brain (Figure 10C). To make network-specific statistical inferences, for each network, we compared FC across each set of connections within the network using a paired t-test; to provide conservative estimates that avoid inflation based on the number of connections, significance was estimated by randomization (10,000 times) for a given pairing of networks. The same procedure was carried out to look at connections between pairings of networks. Bonferroni correction was used to correct for multiple comparisons (Figure 10D).

**Figure 10:**
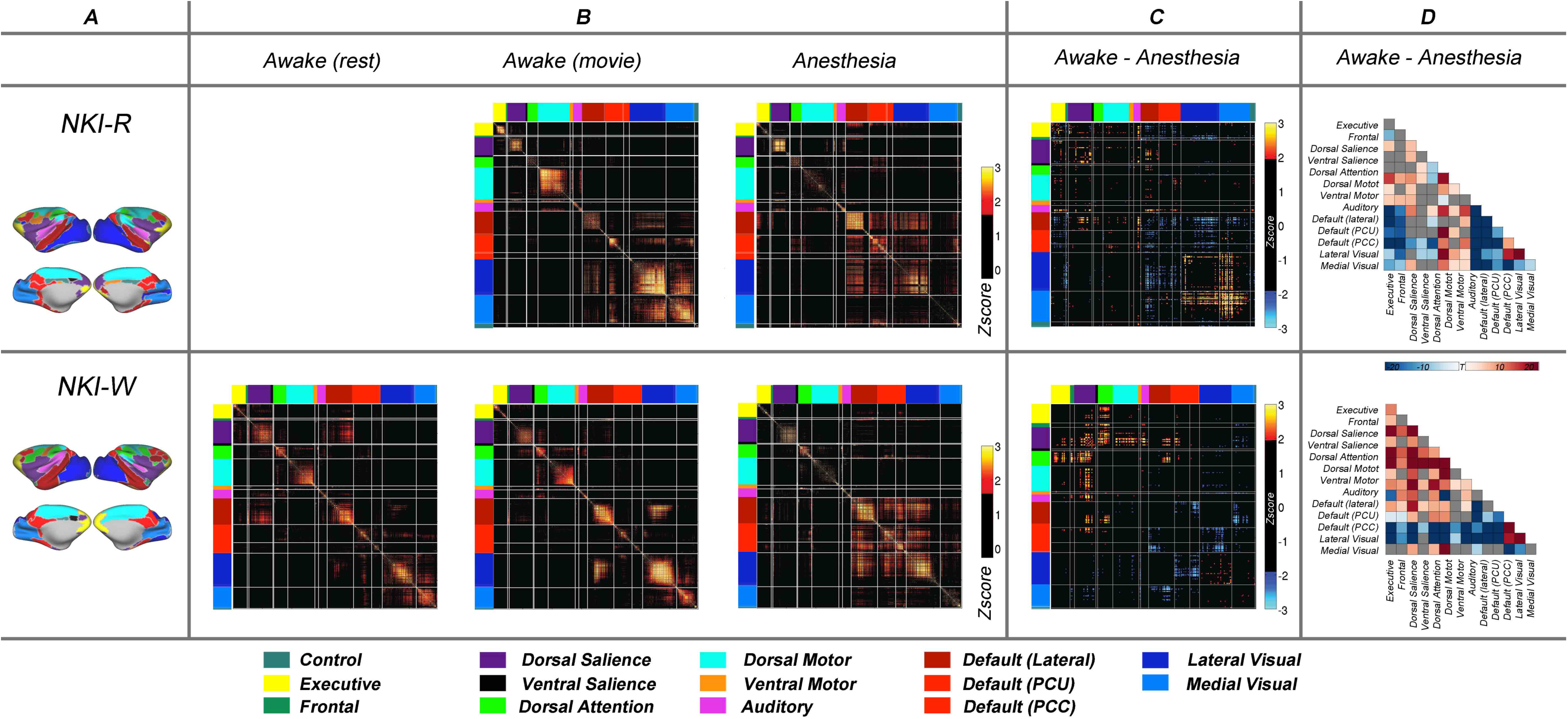
A: The network level of parcellations created by assigning the parcels to the networks defined by group ICA (Ghahremani et al., 2016). B: Functional correlation matrices for the awake states (movie, rest) and anesthesia. To facilitate the visualization, we transformed correlations to z-scores (subtract the mean and divide by standard deviation) and threshold at z-score > 1.645 (p<0.05, one-tail). C: The differences of correlation matrices (r[movie] – r[anesthesia]) that were transformed to z-scores and threshold at z-score > 1.960 (p<0.05, two-tails). D: The significant differences of FC between awake (movie) and anesthesia at network level. The significance was defined by Bonferroni corrected p-score from randomization < 0.05.

Consistent with prior work, we found evidence of anesthesia-related decreases within and between most networks, with specific findings differing somewhat between the two monkeys. Decreases in within-network FC for the dorsal salience, dorsal motor, lateral visual, and medial visual networks were consistent across the two subjects. Of note, although more limited, there was also evidence of anesthesia-related increases in FC - most notably in the posterior cingulate cortex default mode network (PCC-DMN) for the two subjects. Complementing these within-network findings were anesthesia-related increases in between network FC for the visual networks; these increases were most consistent for FC with default network and fronto-parietal components. Of note, while comparison of awake and anesthetized states was not possible for the datasets collected at OHSU, the sparsity of the data collected there was more similar to that obtained under anesthesia at NKI, suggesting some level of consistency.

## DISCUSSION

The present work assessed the ability of a gradient-based FC boundary mapping approach to map the functional organization of the non-human primate brain at the individual level and yield reproducible results. In particular, the present work demonstrated the ability to move beyond network- and region-specific parcellations, to provide full cortical parcellations. Importantly, we found that: 1) parcel boundaries are highly reproducible in individual macaques, though substantial amounts of data are required, 2) resultant parcellations appear to exhibit construct validity per homogeneity analyses and comparison to functional activations in somatosensory cortex, 3) functional connectivity patterns, and their resultant gradients and edge maps appear to be stable across waken states, though markedly different across awake and anesthetized states, 4) enhancement of non-human primate imaging using MION appeared to be essential for obtaining reasonable signal stability in a reasonable time period, and 5) similar to humans, despite gross similarities, individual-specific functional connectivity and parcellation patterns were readily detectable per fingerprinting analyses. As will be discussed in greater detail below, an important reminder that emerged from the present work is that while functional parcellations bear a gross similarity to traditional cytoarchitectonic areas, they are not equivalent.

### Functional Boundaries Are Highly Reproducible in Individual Macaques with Sufficient Data

Our findings regarding reproducibility and data requirements are not surprising given recent work carried out in humans (Gordon et al., 2017a; Laumann et al., 2015; Xu et al., 2016). Similar to the findings from humans, more data is generally better when addressing the reproducible FC (Laumann et al., 2015), though with the largest gains being observed as the data used increases up to ~32 min - a finding that echoes the findings from humans, where the most substantial gains were seen up to around 27–30 minutes. It should be noted that the data needs increase with each additional step of processing (i.e., from FC to gradient and edge calculation). Improvements in reliability for gradients and parcellation slowly but continuously increased as data increased to 40 min, while the largest reproducibility values (spatial correlation and Dice coefficient) were still relatively lower than what can be achieved for FC. These findings suggest that extended data are required to achieve a similar level of reproducibility of parcellations. Additionally, our findings extend recent works emphasizing the importance of contrast agents when scanning non-human primates at 3.0T by demonstrating that the use of MION is a prerequisite to achieving maximal reproducibility for FC, gradients, and functional boundaries maps (Gautama et al., 2003; Grayson et al., 2016; Leite et al., 2002); this is true regardless of whether anesthesia was used or not.

### The Functional Parcellation Show Internal Validity

Directly relevant to the present study, prior studies in humans suggest that the boundary map-based parcellation had highly homogeneous FC patterns at the group and individual level (Gordon et al., 2017a; Laumann et al., 2015; Xu et al., 2016). In the current study, we tested the homogeneity of boundary map-derived parcellations using subsets of the data (subset 1), such that the parcels were generated completely independently from the other subset of the data (subset 2) for each status within each macaque. On average, the parcel homogeneity was above 0.9 with 120 min of awake data. This high degree of homogeneity indicates that most parcels represented areas of similar functional connectivity pattern. It is notable that the homogeneity varied across parcels. As discussed in a previous study (Gordon et al., 2016), the homogeneity of parcels is associated with parcel size, where small parcels are more likely to have higher homogeneity than larger parcels. A few parcels had low homogeneity and some of them, e.g. in the frontal pole, anterior inferior temporal lobe, may be caused by the low-SNR or distortion signal in those areas. Accordingly, it is necessary to use a null model to truly evaluate the homogeneity. By creating the null model from 1000 random rotations of the identical parcellations around the cortical surface, we found that the homogeneities of parcels were significantly higher than the null model in both awake and anesthetized states. Of note, the homogeneity of the cortical parcellations derived from functional data also exhibited a higher degree of homogeneity than those from five established cortical atlases, further increasing confidence in our findings.

A key challenge for any effort focused on cortical parcellation, whether in the human or non-human primate, is determination of the ‘correct’ number of parcels. Regardless of which modality or parcellation method is used, a number of differing criteria can be used to make such determinations. In the current study, the watershed flood algorithm yielded between 110 and 140 parcels per hemisphere, for each macaque. An advantage of this methodology, is the lack of need for a priori specification of the number of parcels to be produced. However, the results can be impacted by properties of the data, such as spatial resolution and smoothness; in this regard, a priori specification of parameters (e.g., number of parcellations) can have advantages. Our estimate for number of parcels converge with current estimates generated from histology-based studies (i.e. cytoarchitecture, myeloarchitecture, and chemoarchitecture), which have increased from 78–88 (Felleman and Van Essen, 1991;

Lewis and Van Essen, 2000; Markov et al., 2010; Van Essen et al., 2012) to 130–160 in each hemisphere (Paxinos and Franklin, 2004; Van Essen et al., 2012). Human work suggested that the number of parcels might increase when we examine gradient across multiple modalities, e.g. area 55b was more distinctive in task activation, and a selective seed connectivity (Glasser et al., 2016). Future work will benefit from integrative examinations across assessment modalities (e.g., functional MRI, diffusion, histology) in the same animals.

### Nonequivalence of functional parcels and traditional cortical areas

It is important to note that functional parcels or “areas” identified in the present work are intended to represent basic units in the macroscale functional brain organization; defined using functional connectivity, they exhibit abrupt changes in connectivity profiles from one parcel to the next (Cohen et al., 2008; Van Essen 2004). Unlike traditionally defined neocortical areas, which are grounded in architectonics, they are not necessarily expected to be stable across all conditions, as functional connectivity patterns are known to vary as a function of state. Previous studies have demonstrated that functional connectivity patterns are sculpted more by structural connectivity under deep anesthesia than the awake state (Barttfeld et al., 2015; Wu et al., 2016). In anesthesia states, when isoflurane levels were varied, the signal fluctuations and connectivity also exhibited distinct profiles at each level within the same animal (Smith et al., 2016; Gao et al., 2016; Peltier et al., 2005; Hutchison et al., 2014). While it is expected that the parcels revealed will in many cases exhibit a gross similarity to traditionally defined neocortical areas, they are not intended to be 1:1. As demonstrated by recent work, the functional areas revealed by gradient-based approaches do appear to exhibit good correspondence with task activations localizers.

A common theme in the parcellation literature is the attempt to establish validity by comparing the parcels obtained with those in atlases derived from more traditional approaches. In this regard, we compared the full brain parcellations derived from our analyses with five atlases, four of which were defined using histological methodologies, and one with multiple modality (e.g., architecture, connections, myeloarchitecture). First, using parcel homogeneity as a metric of fit, we found that the functionally-derived cortical parcellations exhibited superior homogeneity to any of the five atlases. While comparability would be the minimum standard we were aiming for, the superiority was suggestive of three possibilities. Either the atlas is not being optimally registered to each individual, or our functional parcellations successfully accounted for individual-specific variations that cannot be accounted for when applying an atlas, or that functionally-derived parcels are capturing aspects of functional brain organization that are not directly related to traditional cortical areas, or any combination of these (Reveley et al., 2016; Palomero-Gallagher et al., 2016).

Beyond parcel homogeneity, we compared the specific parcels generated from our work with those in the five atlases. While prior work in the human literature has emphasized the ability to recapitulate traditionally defined cortical areas (Gordon et al., 2016; Laumann et al., 2015), the present work raises caveats. While the gross similarity between our parcellations and those of the five atlases was clear, our parcellations did not recapitulates the areas in high detail, regardless of whether one we focused on awake or anesthetized data. This is not necessarily surprising given the many in the literature arguing that structure and function are not 1:1, and showing examples of functional activations failing to follow cortical areas in a precise manner. Intriguingly, for each of the two NKI monkeys, the parcel obtained in the finger area appeared to bound activations observed during somatosensory stimulation – in fact with notably better fit than any of the traditional cortical atlases. As such, we assert the possibility that functionally-derived atlases may be more relevant for functional imaging analyses than traditional, and afford the opportunity to generate individual-specific parcellations in vivo. Further validation using a broader range of mapping approaches is needed to more definitively make this point.

### Rest and Naturalistic Viewing during Awake Imaging Show Similar Functional Properties

Our findings regarding the relative stability of functional connectivity patterns across awake and sensitivities to level of arousal (i.e., sleep, anesthesia) help to synthesize previous findings from the human and non-human primate literatures. Over the past decade, numerous studies have demonstrated the ability to extract highly similar intrinsic functional connectivity networks, regardless of whether imaging is carried out during an active task state or rest (Cole et al., 2014; Fair et al., 2007; Vanderwal et al., 2017). Recent work has increasingly highlighted the potential value of using non-rest states, particularly naturalistic viewing, for assessing functional connectivity, as head motion appears to be lower and tolerability higher. While systematic differences in connectivity patterns are undoubtedly present across states, as revealed by within subject comparisons, differences in connectivity patterns across individuals can be similar - at least for static functional connectivity patterns (Finn et al., 2015; O’Connor et al., 2016; Vanderwal et al., 2017; Wang et al., 2017). Given the various behavioral demands of awake imaging for non-human primate, we have found that naturalistic viewing conditions can be particularly valuable - at least for full-brain parcellations. For those interested in more fine-grained parcellations or temporal dynamics, it is important to further investigate the temporal stability vs. state-dependence of more subtle distinctions within cortical areas (e.g., polar angle and eccentricity mapping in early visual cortex).

### Individual Brain Organization Is Unique While the Awake and Anesthetized States Show Distinct Profiles

As suggested by prior work in humans (Miranda-Domínguez et al, 2014; Xu et al., 2016), we found that individual brain organization can be potentially used for fingerprinting in the monkey. Within the same condition (awake-awake, anesthesia-anesthesia), the within-individual spatial correlations of gradient and edge density maps are explicitly higher than between-individual correlations. Consistent with prior work in human (Miranda-Domínguez et al., 2014; Finn et al., 2015; O’Connor et al., 2016; Vanderwal et al., 2017), the functional brain organization is similar across different awake conditions (i.e., movie, rest), though some decrements in similarities for FC, gradients and edge maps were noted. Importantly, the ability to identify individuals using data collected with anesthesia was notably decreased when attempting to match the same subject across the awake and anesthetized states. This is not surprising, as the similarities for full-brain FC, gradient maps and edge maps were dramatically reduced when looking across the awake and anesthetized states (e.g., ~40% in whole-brain FC similarity). Of note, the current findings were limited in a small sample, the ability to fingerprinting individual brain organization in macaque needs to be confirmed with large datasets in future work.

While the presence of differences in FC properties between awake and anesthetized states was not surprising based on prior work (Hutchison et al., 2013, 2014a; Vincent et al., 2007), some of the specific findings suggest a more nuanced picture than previously appreciated. Consistent with prior studies varying levels of anesthesia, we found evidence of anesthesia-related compromises in within- and between-network connectivity for higher-order associative cortices; the specific networks affected differed somewhat across the two monkeys, with the most consistent findings being observed in the salience and dorsal attention networks. However, unlike prior studies, which largely relied on variations of depth of anesthesia, our comparison of awake states and anesthesia also suggested that the visual networks and their connectivity with default and frontoparietal networks actually showed anesthesia-related increases in connectivity. Overall, these findings echo recent studies of the impact of anesthetics on brain differences in humans, which suggested a loss of complexity in the functional architecture of the brain (Chang et al., 2016; Hutchison et al., 2014b; Peltier et al., 2005; Smith et al., 2016; Wu et al., 2016). Though, further work will be required to rule out other possibilities, such as differences in respiration associated with differing states (awake, anesthesia).

In sum, the present work suggests that each, the awake and the anesthetized states, as well as their differences, are highly stable. Future work would benefit from more detailed examinations of anesthesia effects that include awake imaging, as well as possibly natural sleep. Additionally, an increased focus on potential confounds will be important as the field moves forward; this will require additional monitoring.

### Effects of Sites

Finally, it is worth noting a unique aspect of the present work - the examination of monkey datasets across two institutions. A significant advantage of this strategy was the ability to confirm the reproducibility of findings across two independently collected datasets. Not surprisingly, while we were successful in demonstrating the generality of our findings, we did find evidence of site-related differences. Specifically, we found that the correspondence of gradient and edge maps among subjects was higher within a site than between. Possible factors that might explain site-related variation include differences in: 1) anesthesia protocols (e.g. the knockout agent and delay duration after the time of anesthetics administration); 2) head coils; the OHSU dataset was acquired by a knee coil while NKI dataset was with a surface coil; prior work has demonstrated differences in signal properties (including distortion) of the data obtained from monkeys using these coils (Grayson et al., 2016); 3) rearing histories. Future work would benefit from more coordinated efforts focused on multi-site imaging in the non-human primate as a means of maximizing the reproducibility of findings between laboratories.

### Limitations

It is important to note that despite the various successes of the present work, a number of limitations and potential areas for future optimization exist. As previously discussed, a key limitation of any parcellation methodology is the lack of a gold standard benchmark to help establish validity. The present work made use of comparisons to previously established histologic parcellations to provide insights into validity, as well as the examination of parcel properties (i.e., homogeneity). Future work would benefit from examination of multiple methods of parcellation in the same subjects (e.g., fMRI and diffusion imaging, fMRI and myelin maps, fMRI and histology). Another consideration for future work is examination of the potential impact of preprocessing strategy decisions on the resultant parcels and their stability. Optimization of preprocessing may be particularly useful if focused on bringing down data needs for achieving the same parcellation results. In addition, though the fingerprinting analyses suggest that the functional brain organization is unique and appears to be stable over time, yet the number of samples in the present study was small and large datasets would need to extend and confirm the findings. Finally, the present work relied on the FC similarity measure used in prior studies for gradient-based parcellation. Prior work demonstrated this measure to have favorable properties relative to several other possible measures.

However, an exhaustive examination of measures to be used for gradient-based parcellation has yet to be carried out; this would be of benefit both - to determine if some measures may be able to lead to more fine-grained parcellations, and to find a less computationally costly measure if possible. On a related note, a range of alternatives exist for the final parcellation step in the gradient-based strategy - future work determining optimality is merited.

### Conclusions and Future Directions

In summary, the present work demonstrates the ability to achieve highly reproducible, individual-specific cortical parcellations in the non-human primate. Our findings also highlight the many factors that can contribute to stability, in particular the need for sufficient data and contrast agents. Our findings also emphasized the need for consideration of state (i.e., awake vs. anesthetized) when interpreting findings and their reproducibility across studies, as well as during the study design process. Transformation of our non-human primate findings into a common space demonstrated a future direction that can be further optimized to facilitate comparative and translational studies.

## EXPERIMENTAL PROCEDURES

### Dataset 1, NKI-Macaque

The Macaque dataset 1 consisted of two rhesus monkeys (Macaca mulatta, one male, age 6 years, 6.4 kg, marked as NKI-R; one female, age 7 years, 4.5 kg, marked as NKI-W), which was collected from the Nathan Kline Institute for Psychiatric Research. All methods and procedures were approved by the NKI Institutional Animal Care and Use Committee (IACUC) protocol. The monkeys were previously implanted with an MRI-compatible custom built acrylic head post.

### Data Acquisition

Structural MRI data were obtained with a 3.0 Tesla Siemens Tim Trio scanner with an 8-channel surface coil adapted for monkey head scanning. T1-weighted anatomical images (TE=3.87 ms, TR=2500 ms, TI=1200 ms, flip angle=8 degrees, 0.5 mm isotropic voxel) were collected for each macaque. Each macaque was sedated with an initial dose of atropine (0.05 mg/kg IM), dexdomitor (0.02 mg/kg IM) and ketamine (8 mg/kg IM) intubated, and maintained at 0.75% isoflurane anesthesia for the duration of structural MRI procedures. Functional MRI scans were obtained using a gradient echo EPI sequence (TR=2000 ms, TE=16.6 ms, flip angle=45 degree, 1.5x1.5x2mm voxels, 32 slices, FOV=96 x 96 mm). Seven sessions were acquired for NKI-R (294 min in total) and eleven sessions (668 min) for NKI-W. Each session included 4–7 scans, 8–10 minutes per scan. We collected data under anesthesia and while the monkeys were awake. Monocrystalline iron oxide ferumoxytol (MION) solution was injected at iron doses of 10 mg/kg IV prior to the MRI scanning for all MION sessions. See Table S1 for details of the parameters of the data analyzed. Additionally, we collected somatosensory task (2 sessions for NKI-R and 6 sessions for NKI-W) while the monkeys were awake without MION contrast agent. The air puff was used to stimulate the finger tips. The pseudo-randomized left and right stimulation blocks (20 sec) were separated by 20 sec periods of rest. During the awake scanning, the head motion was controlled using head holder, which was implanted stereotaxically on the cranium of NKI-R and NKI-W under anesthesia before the awake studies.

### Dataset 2, OHSU-Macaque

A second macaque dataset consisted of two male rhesus macaques (Macaca mulatta, one male, age 5 years, 8.6 kg, marked as OHSU-1; one male, age 5 years, 7.6 kg, marked as OHSU-2), which were collected at the Oregon Health and Science University. Animal procedures were in accordance with the National Institutes of Health guidelines on the ethical use of animals and were approved by the Oregon National Primate Research Center (ONPRC) Institutional Animal Care and Use Committee.

### Data Acquisition

Structural MRI data were obtained with a 3.0 Tesla Siemen Tim Trio scanner with a 15-channel knee coil adapted for monkey head scanning. T1-weighted anatomical images (TE=3.33 ms, TR=2600 ms, TI=900 ms, flip angle=8 degrees, 0.5 mm isotropic voxel) were collected for each macaque. Functional MRI scan were obtained using a gradient echo EPI sequence (TR=2070 ms, TE=25 ms, flip angle=90 degrees, 1.5x1.5x1.5 mm voxels, 32 slices, FOV=96 x 96 mm). Each macaque was sedated with an initial dose of ketamine (10 mg/kg) for intubation, and thereafter maintained on <1% isoflurane anesthesia during each scan session. Eight sessions were acquired for each monkey; For each session, 30 min BOLD scan without MION and 30 min scan with MION were acquired at the same day; The BOLD scan started at 45 min after the monkey was anesthetized, followed by the MION scan. The MION solution was injected at iron doses of 8 mg/kg IV. We collected all data at this site under anesthesia. See Table S1 for details of the parameters of the data analyzed.

### Image Preprocessing

Structural image processing included the following steps: 1) spatial noise removal by a non-local mean filtering operation (NKI-dataset) (Zuo and Xing, 2011, 2014), 2) constructing an average structural volume from multiple T1-weight images, 3) brain extraction and tissue segmentation into gray matter (GM), white matter (WM) and cerebrospinal fluid (CSF); this was performed by FSL, FreeSurfer and ANTs, followed by manual editing to fix the mis-segmentations, 4) reconstructing the native white matter and pial surfaces using FreeSurfer, and 5) registering the native surface to a hybrid left-right template surface (Yerkes19 macaque atlas; (Donahue et al., 2016). The Yerkes19 template was created from 19 adult macaques T1 images acquired at the Yerkes Primate Research Center and processed with customized Human Connectome Project (HCP) pipeline (Donahue et al., 2016; Glasser et al., 2013). The left and right hemisphere surfaces were registered with each other and down-sampled from 164k_fs_LR surface to a 10k_fs_LR (10,242 vertices) surface.

Functional image preprocessing included the following steps: 1) discarding of the first five volumes of the time series and compressing temporal spikes (AFNI 3dDespike), 2) slice timing correction, motion correction, and bias field correction (OHSU-dataset), 3) normalizing the 4D global mean intensity, 3) regressing out nuisance signals including the mean time series from WM and CSF masks, the Friston-24 motion parameters (Yan et al., 2013), as well as linear and quadratic trends, 4) band-passed filtering (0.01 < f <0.1Hz) of the residuals to extract the low-frequency fluctuations, 5) registering the functional image to the anatomical space and projecting it onto the native middle surface, 6) spatial smoothing with a 4 mm full width at half maximum kernel along the native surface and down-sampling to the 10k surface.

### Quality Control Procedure

Frame-wise displacement (FD) and mean FD were calculated to quantify the head micro-movements. The mean FD of the scans included in our analysis were less than 0.25 mm during awake scanning and 0.05 mm under anesthesia. The average mean-FD was 0.11 (SD=0.05) across all the awake scans of two macaques and 0.023 (SD=0.006) across all the anesthetized scans of all four of the macaques.

### Surface-based FC Calculation and Gradient-based Parcellation

The cortical surface was reconstructed to provide an accurate representation of morphology and topology of the brain for each macaque. The volumetric fMRI data was aligned to the anatomical space and then projected to the native middle cortical surface. Then the time series were smoothed along the high-resolution (about 164k vertices for each hemisphere) native middle surface (FWHM = 4 mm) and down resampled to a coarser (10,242 vertices for each hemisphere) template surface. Based on the prior study in human work (Xu et al., 2016), the smoothing kernel is chosen about twice that of the surface (10k surface) that the gradients were calculated on.

The procedures of surface-base FC calculation and gradient-based parcellation were similar as previous studies described in Laumann et al. (2015) and Xu et al. (2016). In brief, the functional connectivity (FC) map in full cortex of the gray matter tissue was first computed using the time course for each vertex, resulting in a (10,242 vertices x 20,484 vertices) matrix. The distributions of the resulting correlation values were standardized to the normal distribution using Fisher’s r-to-z transform. Then a FC similarity profile was calculated for each vertex on the surface from the spatial correlation between the vertex’s FC map and the FC map of every other vertex, resulting in a 10,242 vertices x 10,242 vertices symmetric matrix. Each column (or row) of this matrix represents the FC similarity map for each surface vertex. The spatial gradient (i.e., the first spatial derivative) of each FC similarity map was computed on the native middle surface to measure the degree of the transition in FC profile at each vertex, resulting in 10k gradient maps for each hemisphere. A ‘watershed by flooding’ algorithm (Gordon et al., 2016a) was then applied to each of the gradient maps - resulting in 10k binarized edge maps. Finally, the 10k gradient and edge maps were averaged to generate the final gradient and an edge density map. By applying the same watershed algorithm to the edge density map to yield the final parcel map.

### Parcellation Validation

For each Macaque, in order to compare the homogeneity across different conditions (awake vs. anesthesia, MION vs. no MION), the parcel homogeneity was evaluated using Kendall’ coefficient at all vertices in the parcel for each condition (Xu et al., 2016). Consistent with previous human work, the overall homogeneity of the parcellation was compared to a null model created from 1000 random rotations of the parcellations on the cortical surface (Gordon et al., 2016).

### Evaluating the Stability of FC, Gradients, and Edge Density

For each macaque, we randomly split the data from all sessions into halves within each condition, setting aside one half of the data as a reference (subset 2). We then randomly selected samples from subset 1 of 8, 12, 16, 20, 24, 28, 32, 40, 48, 56, 72, and 88 min (up to the half of the sample for each macaque in each condition) to examine convergent estimates of FC, gradient and edges. Of note, previous findings suggested that shorter sampling of contiguous data over more sessions can facilitate the convergence of stability. Hence the subsamples of 8 to 88 min data were generated from multiple 2 min contiguous segments from the subset 1. Parcel-based FC was calculated to evaluate the areal-level cortical network and permuted 1000 times at each time point. Gradient and edge density maps were also calculated to examine the convergence of the parcellation. To facilitate the computational efficiency for gradient and edge density calculations, the results of gradient and edge density were performed on the down-resampled 400 regions of vertices of interest for 100 times at each time point.

### Network Assignment

To characterize a large-scale system of parcellations created in the current study, we assigned a network identity to parcels using the previously established network definition from Ghahremani et al. (2016) The matching procedure was similar with Gordon et al. (2016). Specifically, we extracted the time series and averaged it within each parcel. The averaged time series were then correlated against all other times series across the cortical surface to obtain a connectivity map. Connections were excluded if the geodesic distance between parcel centers was less than 10 mm to remove the purely local connectivities (Power et al., 2011) from spatial smoothing. After that, we thresholded and binarized the connectivity map at the top 5% of connectivity strengths. This resulted in a binarized map of regions with high connectivity between parcels. Then we examined the overlap of this binarized map to the binarized group ICA Z-map (Z>2.33, p<0.001) from Ghahremani et al. (2016) and defined the best match using the Dice coefficient. Then neighboring parcels assigned to the same network were merged to create a final network-patch.

## AUTHOR CONTRIBUTIONS

T.X., C.S., D.F., and M.P.M. designed the study. T.X. analyzed the data with support from J.R., E.F., and D.S. A.F., G.L., D.R., J.B., E.E., O.MD. and A.P performed the experiments and provided support. T.X. M.P.M wrote the manuscript with contributions from E.L.S., C.S. and D.F. All authors commented on the manuscript.

## REFERENCE

Brodmann, K. (1909). Vergleichende Lokalisationslehre der Grosshirnrinde in ihren Prinzipien dargestellt auf Grund des Zellenbaues.

Buckner, R.L., and Yeo, B.T.T. (2014). Borders, map clusters, and supra-areal organization in visual cortex. Neuroimage 93 Pt 2, 292–297.

Barttfeld, Pablo, Lynn Uhrig, Jacobo D Sitt, Mariano Sigman, and Béchir Jarraya. (2015). Signature of Consciousness in the Dynamics of Resting-State Brain Activity. Proc. Natl. Acad. Sci. U. S. A. 112 (37): E5219–20.

Cerliani, L., D’Arceuil H., and Thiebaut de Schotten, M. (2017). Connectivity-based parcellation of the macaque frontal cortex, and its relation with the cytoarchitectonic distribution described in current atlases. Brain Struct. Funct. 222, 1331–1349.

Chang, C., Leopold, D.A., Schölvinck, M.L., Mandelkow, H., Picchioni, D., Liu, X., Ye, F.Q., Turchi, J.N., and Duyn, J.H. (2016). Tracking brain arousal fluctuations with fMRI. Proc. Natl. Acad. Sci. U. S. A. 113, 4518–4523.

Cohen, A.L., Fair, D.A., Dosenbach, N.U.F., Miezin, F.M., Dierker, D., Van Essen, D.C., Schlaggar, B.L., and Petersen, S.E. (2008). Defining functional areas in individual human brains using resting functional connectivity MRI. Neuroimage 41, 45–57.

Cole, M.W., Bassett, D.S., Power, J.D., Braver, T.S., and Petersen, S.E. (2014). Intrinsic and task-evoked network architectures of the human brain. Neuron 83, 238–251.

Craddock, R.C., Cameron Craddock, R., James, G.A., Holtzheimer, P.E., Hu, X.P., and Mayberg, H.S. (2011). A whole brain fMRI atlas generated via spatially constrained spectral clustering. Hum. Brain Mapp. 33, 1914–1928.

Donahue, C.J., Sotiropoulos, S.N., Jbabdi, S., Hernandez-Fernandez M., Behrens, T.E., Dyrby, T.B., Coalson, T., Kennedy, H., Knoblauch, K., Van Essen, D.C., et al. (2016). Using Diffusion Tractography to Predict Cortical Connection Strength and Distance: A Quantitative Comparison with Tracers in the Monkey. J. Neurosci. 36, 6758–6770.

Essen, D.V. (2012). Surface-based analyses of human, macaque, and chimpanzee cortical organization. J. Vis. 12, 1377–1377.

Fair, D.A., Schlaggar, B.L., Cohen, A.L., Miezin, F.M., Dosenbach, N.U.F., Wenger, K.K., Fox, M.D., Snyder, A.Z., Raichle, M.E., and Petersen, S.E. (2007). A method for using blocked and event-related fMRI data to study “resting state” functional connectivity. Neuroimage 35, 396–405.

Felleman, D.J., and Van Essen, D.C. (1991). Distributed hierarchical processing in the primate cerebral cortex. Cereb. Cortex 1, 1–47.

Finn, E.S., Shen, X., Scheinost, D., Rosenberg, M.D., Huang, J., Chun, M.M., Papademetris, X., and Constable, R.T. (2015). Functional connectome fingerprinting: identifying individuals using patterns of brain connectivity. Nat. Neurosci. 18, 1664–1671.

Gao, Yu Rong, Yuncong Ma, Qingguang Zhang, Winder Aaron T., Zhifeng Liang, Lilith Antinori, Patrick J. Drew, and Nanyin Zhang. (2016) Time to Wake up: Studying Neurovascular Coupling and Brain-Wide Circuit Function in the Un-Anesthetized Animal. NeuroImage 153, 382–398.

Gautama, T., Mandic, D.P., and Van Hulle, M.M. (2003). Signal nonlinearity in fMRI: a comparison between BOLD and MION. IEEE Trans. Med. Imaging 22, 636–644.

Ghahremani, M., Matthew Hutchison, R., Menon, R.S., and Everling, S. (2016). Frontoparietal Functional Connectivity in the Common Marmoset. Cereb. Cortex.

Glasser, M.F., Sotiropoulos, S.N., Wilson, J.A., Coalson, T.S., Fischl, B., Andersson, J.L., Xu, J., Jbabdi, S., Webster, M., Polimeni, J.R., et al. (2013). The minimal preprocessing pipelines for the Human Connectome Project. Neuroimage 80, 105–124.

Glasser, M.F., Coalson, T.S., Robinson, E.C., Hacker, C.D., Harwell, J., Yacoub, E., Ugurbil, K., Andersson, J., Beckmann, C.F., Jenkinson, M., et al. (2016). A multi-modal parcellation of human cerebral cortex. Nature 536, 171–178.

Gordon, E.M., Laumann, T.O., Adeyemo, B., Huckins, J.F., Kelley, W.M., and Petersen, S.E. (2016). Generation and Evaluation of a Cortical Area Parcellation from Resting-State Correlations. Cereb. Cortex 26, 288–303.

Gordon, E.M., Laumann, T.O., Adeyemo, B., and Petersen, S.E. (2017a). Individual Variability of the System-Level Organization of the Human Brain. Cereb. Cortex 27, 386–399.

Gordon, E.M., Laumann, T.O., Adeyemo, B., Gilmore, A.W., Nelson, S.M., Dosenbach, N.U.F., and Petersen, S.E. (2017b). Individual-specific features of brain systems identified with resting state functional correlations. Neuroimage 146, 918–939.

Grayson, D.S., Bliss-Moreau E., Machado, C.J., Bennett, J., Shen, K., Grant, K.A., Fair, D.A., and Amaral, D.G. (2016). The Rhesus Monkey Connectome Predicts Disrupted Functional Networks Resulting from Pharmacogenetic Inactivation of the Amygdala. Neuron 91, 453–466.

Hutchison, R.M., and Everling, S. (2012). Monkey in the middle: why non-human primates are needed to bridge the gap in resting-state investigations. Front. Neuroanat. 6, 29.

Hutchison, R.M., and Everling, S. (2014). Broad intrinsic functional connectivity boundaries of the macaque prefrontal cortex. Neuroimage 88, 202–211.

Hutchison, R.M., Gallivan, J.P., Culham, J.C., Gati, J.S., Menon, R.S., and Everling, S. (2012a). Functional connectivity of the frontal eye fields in humans and macaque monkeys investigated with resting-state fMRI. J. Neurophysiol. 107, 2463–2474.

Hutchison, R.M., Womelsdorf, T., Gati, J.S., Leung, L.S., Menon, R.S., and Everling, S. (2012b). Resting-State Connectivity Identifies Distinct Functional Networks in Macaque Cingulate Cortex. Cereb. Cortex 22, 1294–1308.

Hutchison, R.M., Womelsdorf, T., Gati, J.S., Everling, S., and Menon, R.S. (2013). Resting-state networks show dynamic functional connectivity in awake humans and anesthetized macaques. Hum. Brain Mapp. 34, 2154–2177.

Hutchison, R.M., Hutchison, M., Manning, K.Y., Menon, R.S., and Everling, S. (2014a). Isoflurane induces dose-dependent alterations in the cortical connectivity profiles and dynamic properties of the brain’s functional architecture. Hum. Brain Mapp. 35, 5754–5775.

Hutchison, R.M., Hutchison, M., Manning, K.Y., Menon, R.S., and Everling, S. (2014b). Isoflurane induces dose-dependent alterations in the cortical connectivity profiles and dynamic properties of the brain’s functional architecture. Hum. Brain Mapp. 35, 5754–5775.

Hutchison, R.M., Culham, J.C., Flanagan, J.R., Everling, S., and Gallivan, J.P. (2015). Functional subdivisions of medial parieto-occipital cortex in humans and nonhuman primates using resting-state fMRI. Neuroimage 116, 10–29.

Laumann, T.O., Gordon, E.M., Adeyemo, B., Snyder, A.Z., Joo, S.J., Chen, M.-Y., Gilmore, A.W., McDermott K.B., Nelson, S.M., Dosenbach, N.U.F., et al. (2015a). Functional System and Areal Organization of a Highly Sampled Individual Human Brain. Neuron 87, 657–670.

Leite, F.P., Tsao, D., Vanduffel, W., Fize, D., Sasaki, Y., Wald, L.L., Dale, A.M., Kwong, K.K., Orban, G.A., Rosen, B.R., et al. (2002). Repeated fMRI using iron oxide contrast agent in awake, behaving macaques at 3 Tesla. Neuroimage 16, 283–294.

Lewis, J.W., and Van Essen, D.C. (2000). Mapping of architectonic subdivisions in the macaque monkey, with emphasis on parieto-occipital cortex. J. Comp. Neurol. 428, 79–111.

Markov, N.T., Misery, P., Falchier, A., Lamy, C., Vezoli, J., Quilodran, R., Gariel, M.A., Giroud, P., Ercsey-Ravasz M., Pilaz, L.J., et al. (2010). Weight Consistency Specifies Regularities of Macaque Cortical Networks. Cereb. Cortex 21, 1254–1272.

Mars, R.B., Jbabdi, S., Sallet, J., O’Reilly J.X., Croxson, P.L., Olivier, E., Noonan, M.P., Bergmann, C., Mitchell, A.S., Baxter, M.G., et al. (2011). Diffusion-weighted imaging tractography-based parcellation of the human parietal cortex and comparison with human and macaque resting-state functional connectivity. J. Neurosci. 31, 4087–4100.

Miranda-Domínguez, O., Mills, B.D., Grayson, D., Woodall, A., Grant, K.A., Kroenke, C.D., and Fair, D.A. (2014). Bridging the gap between the human and macaque connectome: a quantitative comparison of global interspecies structure-function relationships and network topology. J. Neurosci. 34, 5552–5563.

Nelson, E.E., and Winslow, J.T. (2008). Non-Human Primates: Model Animals for Developmental Psychopathology. Neuropsychopharmacology 34, 90–105.

Neubert, F.-X., Mars, R.B., Sallet, J., and Rushworth, M.F.S. (2015). Connectivity reveals relationship of brain areas for reward-guided learning and decision making in human and monkey frontal cortex. Proc. Natl. Acad. Sci. U. S. A. 112, E2695–E2704.

O’Connor D., Potler, N.V., Kovacs, M., Xu, T., Ai, L., Pellman, J., Vanderwal, T., Parra, L., Cohen, S., Ghosh, S., et al. (2016). The Healthy Brain Network Serial Scanning Initiative: A resource for evaluating inter-individual differences and their reliabilities across scan conditions and sessions.

Palomero-Gallagher, Nicola, Simon B Eickhoff, Felix Hoffstaedter, Axel Schleicher, Hartmut Mohlberg, Brent A Vogt, Katrin Amunts, and Karl Zilles. (2015). Functional Organization of Human Subgenual Cortical Areas: Relationship between Architectonical Segregation and Connectional Heterogeneity. NeuroImage 115, 177–190.

Paxinos, G., and Franklin, K.B.J. (2004). The Mouse Brain in Stereotaxic Coordinates (Gulf Professional Publishing).

Peltier, S.J., Kerssens, C., Hamann, S.B., Sebel, P.S., Byas-Smith M., and Hu, X. (2005). Functional connectivity changes with concentration of sevoflurane anesthesia. Neuroreport 16, 285–288.

Phillips, K.A., Bales, K.L., Capitanio, J.P., Conley, A., Czoty, P.W., ‘t Hart, B.A., Hopkins, W.D., Hu, S.-L., Miller, L.A., Nader, M.A., et al. (2014). Why Primate Models Matter. Am. J. Primatol. 76, 801.

Power, J.D., Cohen, A.L., Nelson, S.M., Wig, G.S., Barnes, K.A., Church, J.A., Vogel, A.C., Laumann, T.O., Miezin, F.M., Schlaggar, B.L., et al. (2011). Functional network organization of the human brain. Neuron 72, 665–678.

Reveley, Colin, Audrūnas Gruslys, Frank Q Ye, Daniel Glen, Jason Samaha, Brian E Russ, Ziad Saad, Anil K Seth, David A Leopold, and Kadharbatcha S Saleem. (2017). Three-Dimensional Digital Template Atlas of the Macaque Brain. Cerebral Cortex. 27(9), 4463–4477.

Rilling, J.K. (2014). Comparative primate neuroimaging: insights into human brain evolution.

Schaefer, A., Kong, R., Gordon, E.M., Laumann, T.O., Zuo, X.-N., Holmes, A., Eickhoff, S.B., and Thomas Yeo, B.T. (2017). Local-Global Parcellation of the Human Cerebral Cortex From Intrinsic Functional Connectivity MRI.

Schönwiesner, M., Dechent, P., Voit, D., Petkov, C.I., and Krumbholz, K. (2014). Parcellation of Human and Monkey Core Auditory Cortex with fMRI Pattern Classification and Objective Detection of Tonotopic Gradient Reversals. Cereb. Cortex 25, 3278–3289.

Shen, K., Bezgin, G., Hutchison, R.M., Gati, J.S., Menon, R.S., Everling, S., and McIntosh, A.R. (2012). Information processing architecture of functionally defined clusters in the macaque cortex. J. Neurosci. 32, 17465–17476.

Smith, J.B., Liang, Z., Watson, G.D.R., Alloway, K.D., and Zhang, N. (2016). Interhemispheric resting-state functional connectivity of the claustrum in the awake and anesthetized states. Brain Struct. Funct.

Vanderwal, T., Eilbott, J., Finn, E.S., Craddock, R.C., Turnbull, A., and Castellanos, F.X. (2017a). Individual differences in functional connectivity during naturalistic viewing conditions. Neuroimage.

Vanduffel, W., Zhu, Q., and Orban, G.A. (2014a\). Monkey cortex through fMRI glasses. Neuron 83, 533–550.

Van Essen, D.C., and Dierker, D.L. (2007). Surface-based and probabilistic atlases of primate cerebral cortex. Neuron 56, 209–225.

Van Essen, D.C., Glasser, M.F., Dierker, D.L., and Harwell, J. (2012). Cortical parcellations of the macaque monkey analyzed on surface-based atlases. Cereb. Cortex 22, 2227–2240.

Van Essen, D.C., Donahue, C., Dierker, D.L., and Glasser, M.F. (2016). Parcellations and Connectivity Patterns in Human and Macaque Cerebral Cortex. In Micro-, Meso- and Macro-Connectomics of the Brain, H. Kennedy, D.C. Van Essen, and Y. Christen, eds. (Cham (CH): Springer),.

Vincent, J.L., Patel, G.H., Fox, M.D., Snyder, A.Z., Baker, J.T., Van Essen, D.C., Zempel, J.M., Snyder, L.H., Corbetta, M., and Raichle, M.E. (2007). Intrinsic functional architecture in the anaesthetized monkey brain. Nature 447, 83–86.

Wang, J., Ren, Y., Hu, X., Nguyen, V.T., Guo, L., Han, J., and Guo, C.C. (2017). Testretest reliability of functional connectivity networks during naturalistic fMRI paradigms.

Hum. Brain Mapp. 38, 2226–2241.

Wig, G.S., Laumann, T.O., and Petersen, S.E. (2014). An approach for parcellating human cortical areas using resting-state correlations. Neuroimage 93 Pt 2, 276–291.

Wu, T.-L., Mishra, A., Wang, F., Yang, P.-F., Gore, J.C., and Chen, L.M. (2016). Effects of isoflurane anesthesia on resting-state fMRI signals and functional connectivity within primary somatosensory cortex of monkeys. Brain Behav. 6, e00591.

Xu, T., Opitz, A., Craddock, R.C., Wright, M.J., Zuo, X.-N., and Milham, M.P. (2016). Assessing Variations in Areal Organization for the Intrinsic Brain: From Fingerprints to Reliability. Cereb. Cortex. 26, 4192–4211

Yan, C.-G., Craddock, R.C., He, Y., and Milham, M.P. (2013). Addressing head motion dependencies for small-world topologies in functional connectomics. Front. Hum. Neurosci. 7, 910.

Zuo, X.-N., and Xing, X.-X. (2011). Effects of non-local diffusion on structural MRI preprocessing and default network mapping: statistical comparisons with isotropic/anisotropic diffusion. PLoS One 6, e26703.

Zuo, X.-N., and Xing, X.-X. (2014). Test-retest reliabilities of resting-state FMRI measurements in human brain functional connectomics: a systems neuroscience perspective. Neurosci. Biobehav. Rev. 45, 100–118.

